# UPF3A is a ubiquitously expressed NMD factor among mouse tissues

**DOI:** 10.1101/2023.02.06.526166

**Authors:** Xin Ma, Yan Li, Chengyan Chen, Tangliang Li

## Abstract

Nonsense-mediated mRNA decay (NMD), an important post-transcriptional regulatory mechanism in gene expression, is actively involved in a series of cellular and physiological processes, thus controlling cell fate and tissue homeostasis. Defects in NMD cause human diseases such as neurodevelopmental disorders, tumorigenesis and autoimmunity. UPF3 (Up-frameshift protein 3), first identified in the baker’s yeast, is a core NMD factor. UPF3A and UPF3B, the two UPF3 paralogs emerging in vertebrates, have either activating or suppressing roles in NMD. Previous studies found that UPF3B protein is ubiquitously expressed in almost all mammalian organs, while UPF3A protein is hardly detectable in most of mammalian tissues, except in the testis. One hypothesis explaining this phenomena is the functional antognism between UPF3A and UPF3B in NMD. Thus, UPF3B competitively binds to UPF2 with higher affinity than UPF3A, which finally destabilizes UPF3A protein. In the present study, we quantitatively evaluated the expression of UPF3A and UPF3B in nine major tissues and reproductive organs of wild type male and female mice. Our study confirmed that UPF3A has the highest expression in male germlines. To our surprise, we found in most tissues, including brain and thymus, the protein level of UPF3A is comparable with that of UPF3B. In spleen and lung, UPF3A is higher than UPF3B. These findings are further supported by publicly available gene expression data. Thus, our study demonstrated that UPF3A protein is ubiquitously expressed in mouse tissues, and may play important roles in the homeostasis of multiple mammalian tissues.

## 1 Induction

Nonsense-mediated mRNA decay (NMD) is a highly conserved post-transcriptional gene regulatory mechanism in eukaryotes (1-6). NMD is considered as an RNA surveillance machinery, which is an essential biological pathway to regulate mRNA degradation and eventual protein expression. NMD can recognize and degrade aberrant mRNAs, which contain premature termination codons (PTCs) located at least 50∼55 nucleotides upstream of the last exon-exon junction, to prevent possible toxic effects from truncated proteins translated (1-3, 5, 6). Furthermore, recent studies show that NMD can recognize the special structures of mRNAs with physiological functions, such as long 3′ untranslated region (3’ UTR) and upstream open reading frames of mRNAs, so as to regulate cell fate (7). Thus, NMD is essential for embryonic development and tissue homeostasis in mammals (2, 8-12).

NMD was first discovered in *Saccharomyces cerevisiae* (13). Its core factors include up-frameshift protein (UPF)-1, UPF2 and UPF3 identified in the unicellular eukaryotic organism (yeast) (13, 14), and suppressors with morphological effects on genitalia (SMG)-5, SMG6, and SMG7 in multicellular organisms (*worms, frogs and mammals*), which provide additional layers of regulations on NMD execution (5, 15-17). Among these NMD core factors, UPF3 is a mysterious one. There is only a single UPF3 in invertebrate organisms, including yeasts, worms and flies, while UPF3A and UPF3B, two paralogs of UPF3, are proposed to be generated by a gene duplication event during the emergence of vertebrate animals (18, 19). In mouse and human, UPF3B is considered as the dominant UPF3 paralog expressed in almost all tissues, expect that UPF3A is highly expressed in male germlines (18, 20, 21). In human, Serin *et al.* and Tarpey *et al.* used the Southern blotting method and found that *UPF3B* transcipts are highly produced in all major ograns including bone marrow, brain, lung, testis and intenstine, etc., while *UPF3A* transcripts are highly enriched in human testis. Only little amounts of *UPF3A* mRNAs are found in human bone marrow, brain, intestine samples (20, 21). In mouse, Upf3a mRNAs in testis is around 40-folds higher than other organs. In consistent to the RNA data in human and mouse, Shum *et al.* used antibody based-apprach and found UPF3A protein is almost undetectable in the brain, heart, kidney, liver and spleen of adult mice, while UPF3B is abundantly expressed in these tissues (18). The “under-represented” UPF3A protein in most mammalian tissues was patially explained by the hypothesis that UPF3B competes over UPF3A on binding to NMD factor UPF2, which destabilizes UPF3A protein (18, 22). In accordance with this hypothesis, the depletion of UPF3B protein does result in a dramatic increase of UPF3A protein (18, 22, 23). Thus, protein expressions of UPF3A and UPF3B could be mutually exclusive in most mammalian organs, which supports the functional antognism between UPF3A and UPF3B.

Another mystery on UPF3 paralogs is their biological functions in activating or suppressing NMD. UPF3B is critical during the mammalian neurodevelopment (19, 21, 24, 25). UPF3B mutations cause schizophrenia, intellectual disability and autism spectrum disorders in humans (21, 24). Thus, the biology of UPF3B is better characterized in mammals (19). Recent findings reconcile a conclusion that UPF3B is not only a weak NMD activator (16, 20, 26-28), but also is involved in the regulation of translation termination (27, 29). UPF3A, when tethered to mRNAs at a site downstream of a stop codon, elicits the NMD (26, 27). Furthermore, UPF3A knockdown weakly stabilizes a subset of NMD targets (22). Thus, UPF3A, as UPF3B, could positively regulate NMD. Intriguingly, Shum *et al.* reported that downregulation of UPF3A leads to increased NMD activity in human and mouse cells, and overexpression of UPF3A results in NMD inhibition (18). These results imply that UPF3A is an NMD repressor (18). Two more recent studies revisited the interrelationship of UPF3 paralogs in several human cell lines. They found that knockout of UPF3A or UPF3B alone weakly inhibits NMD, while simultaneous knockout of UPF3A and UPF3B result in significant inhibition of NMD and the strong accumulation of PTC containing mRNAs (16, 28). Thus, UPF3A remains as a weak activator of NMD in human (16, 28).

To further investigate the role of UPF3A in murine NMD, we recently constructed a *Upf3a* conditional knockout mouse line and generated a batch of *Upf3a* inducible knockout embryonic stem cells and rib muscle derived fibroblasts. We found that UPF3A loss does not inhibit NMD in mouse pluripotent and somatic cells (30), supporting the conclusion that UPF3A is not a NMD repressor, but a weak activator (16, 28). Surprisingly, we found that UPF3A, as UPF3B, is adequately expressed in all the mouse cell lines we established. This finding motivated us to re-investigate the physiological expression patterns of UPF3A and UPF3B in different tissues from wild type mice, which stands as the basics in UPF3A and UPF3B biology. We measured the expression of UPF3A and UPF3B at mRNAs and protein levels in nine major mouse tissues (liver, heart, spleen, lung, kidney, thymus, small intestine, cerebral cortex, and olfactory bulb) and reproductive tissues (testis and ovary) from males and females. We found that UPF3A and UPF3B are not mutual-exclusively but adequately and ubiquitously expressed in most of the tissue we investigated. Thus, our finding revises the previous long-standing assumption on UPF3A and UPF3B expression among tissues, and will serve as a reference for further in-depth study on the biology of UPF3 paralogs in mammals.

## 2 Materials and Methods

### 2.1 Mice

Male and female C57BL/6 mice aged 8-10 weeks were maintained under specific pathogen-free conditions at the animal facility of Shandong University (Qingdao, China). All mice had free access to water and food. All animal care and experiments were performed according to the guidelines of the ethics committee (License number: SYDWLL-2022-083).

### 2.2 Real-time quantitative PCR (RT-qPCR)

Total RNAs of mouse tissues were extracted by TRIzol reagent (Takara Bio, Tokyo, Japan), then were used for cDNA synthesis with HiScript^®^ II Q Select RT SuperMix for qPCR (Vazyme Biotech Co., Ltd., Nanjing, China). RT-qPCR was performed on the CFX Real-time PCR system (Bio-Rad, Hercules, CA, USA) using SYBR Green Master qPCR Mix (Tsingke Biotechnology Co., Ltd., Beijing, China). The primers used are listed as bellow: *mUpf3a*: F, GCGCACGATTACTTCGAGGT; R, TCAAAACGGTCTCTGAACAGC; *mUpf3b*: F, AGGAGAAACGAGTGACCCTGT; R, CCTGTTGCGATCCTGCCTA; *mβ-actin*: F, AGAGGGAAATCGTGCGTGAC; R, CAATAGTGATGACCTGGCCGT.

### 2.3 Western blotting

Western blotting analysis followed a previously published method (31). Briefly, RIPA buffer (Radioimmunoprecipitation assay buffer) supplemented with protease/phosphatase inhibitors (APExBIO, Houston, TX, USA) was used for total protein extraction from mouse tissues. The Pierce BCA Protein Assay Kit (Thermo Fisher Scientific, Waltham, MA, USA) was used for measuring the protein concentrations. Sodium dodecyl sulfate-polyacrylamide gel electrophoresis was performed to separate the proteins, which were further transferred to PVDF membranes (Merck Millipore Ltd., Darmstadt, Germany). After blocking with 5% milk-Tris-buffered saline with Tween 20 (TBST) for 2 h, the membranes were incubated with rabbit anti- UPF3A+3B antibody (1:1000, Abcam, Cambridge, UK) or mouse anti-GAPDH antibody (1:5000, Proteintech, Tokyo, Japan) overnight at 4℃, then washed three times with TBST and incubated with corresponding secondary antibodies for 1 h at room temperature. Protein bands were visualized using BeyoECL Plus (Beyotime Biotechnology, Shanghai, China), and quantified using the ImageJ software (National Institutes of Health, Bethesda, MD, USA).

### 2.4 Statistical analysis

Data were statistically analyzed by SPSS (v26.0) software, and all data are presented as means ± standard deviation (SD). *P* value < 0.05 was considered significant. Kolmogorov-Smirnov test was used to determine whether the data followed the normal distribution. Comparisons between the two groups were executed by unpaired Student’s *t*-test or Mann-Whitney *U*-test.

## 3 Results

### 3.1 mRNA expression of *Upf3a* and *Upf3b* among different tissues in adult mice

We examined the mRNA expression of *Upf3a* and *Upf3b* in nine major organs (liver, heart, spleen, lung, kidney, thymus, small intestine, cerebral cortex, and olfactory bulb), and reproductive tissues (testis and ovary) of adult C57BL/6 mice. RT-qPCR analysis showed that small intestine has the lowest expression of *Upf3a* and *Upf3b* in both sexes (Figure 1A and 1B). In adult males, the mRNA level of *Upf3a* is the highest in the testis, 20 times more than that in the liver (Figure 1A). In hematopoietic organs, including spleen and thymus, mRNAs of *Upf3a* and *Upf3b* are generally lower than those in liver, heart, lung, kidney, cerebral cortex and olfactory bulb from males. In females, no significant difference in the mRNA expression of *Upf3a* and *Upf3b* is found between tissues (Figure 1B), although we could still notice that hematopoietic organs have lower expression of *Upf3a* and *Upf3b*.

**Figure 1.**
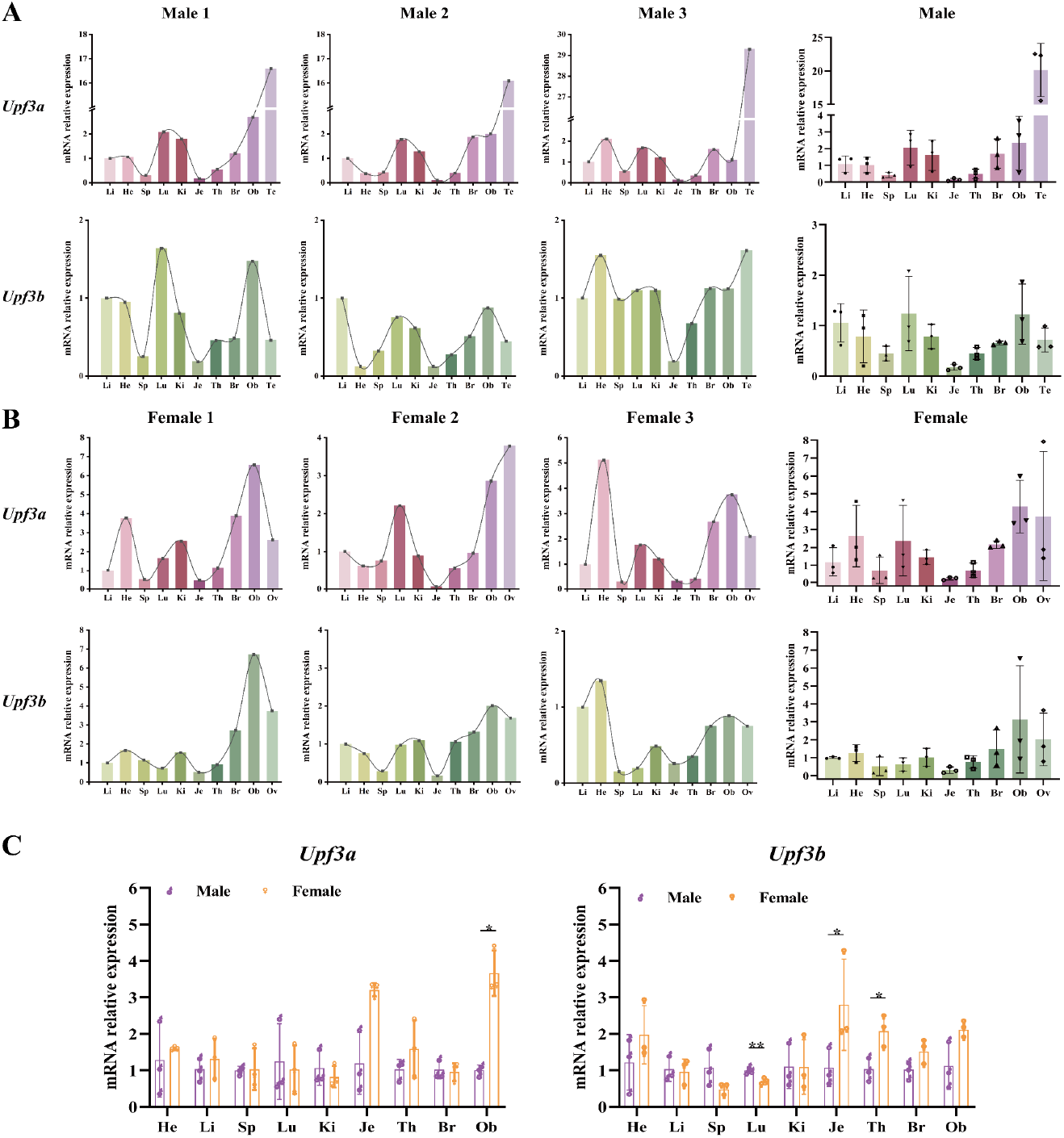
The mRNA relative expression of *Upf3a* and *Upf3b* in tissues of C57BL/6 mice measured by Real-time quantitative PCR. **(A)** Male mice. **(B)** Female mice. **(C)** Comparison of male and female mice. n=3. Results are represented as mean ± SD. *, P < 0.05; **, P < 0.01. He: heart; Li: Liver; Sp: spleen; Lu: lung; Ki: kidney; Je: jejunum; Th: thymus; Br: brain; Ob: olfactory bulb; Te: testis; Ov: ovary.

Next, we compared the differences in the mRNA expression of *Upf3a* and *Upf3b* between the sexes. In general, *Upf3a* and *Upf3b* mRNA expressions are comparable between sexes. However, *Upf3a* mRNA expression in the olfactory bulbs from females, and *Upf3b* mRNA expression in small intestines and thymuses of females are significantly higher than those of male mice, while the mRNA expression of *Upf3b* in the lungs of males is higher than that in females (Figure 1C).

### 3.2 Protein expression of UPF3A and UPF3B among different tissues in adult mice

RT-qPCR analysis with a different set of primers does not allow for the direct comparison of the expression of two genes. Furthermore, antibody-based approach normally could distinguish the relative expression of protein isoforms of a single gene, and could not determine the relative expression of two gene products in a single blot. We previously characterized a commercial antibody which has cross-reactivities with mouse UPF3A and UPF3B proteins. This antibody allows us to directly compare the relative expression of UPF3A and UPF3B in one immunological reaction. We first explored the tissue specific expression of UPF3A and UPF3B in male and female mice, respectively.

As expected, we found UPF3B is ubiquitously expressed in most of mouse tissues, including the liver, spleen, lung, kidney, thymus, cerebral cortex, and olfactory bulb from both sexes and ovaries in females (Figure 2). However, hearts and small intestines from both sexes have almost negligible expression of UPF3B. For UPF3A, as expected, it is highly expressed in male testis. Interestingly, we found UPF3A is evidently expressed in almost all the mouse tissues, including the liver, spleen, lung, thymus, cerebral cortex, and olfactory bulb from both sexes and ovaries in females.

**Figure 2.**
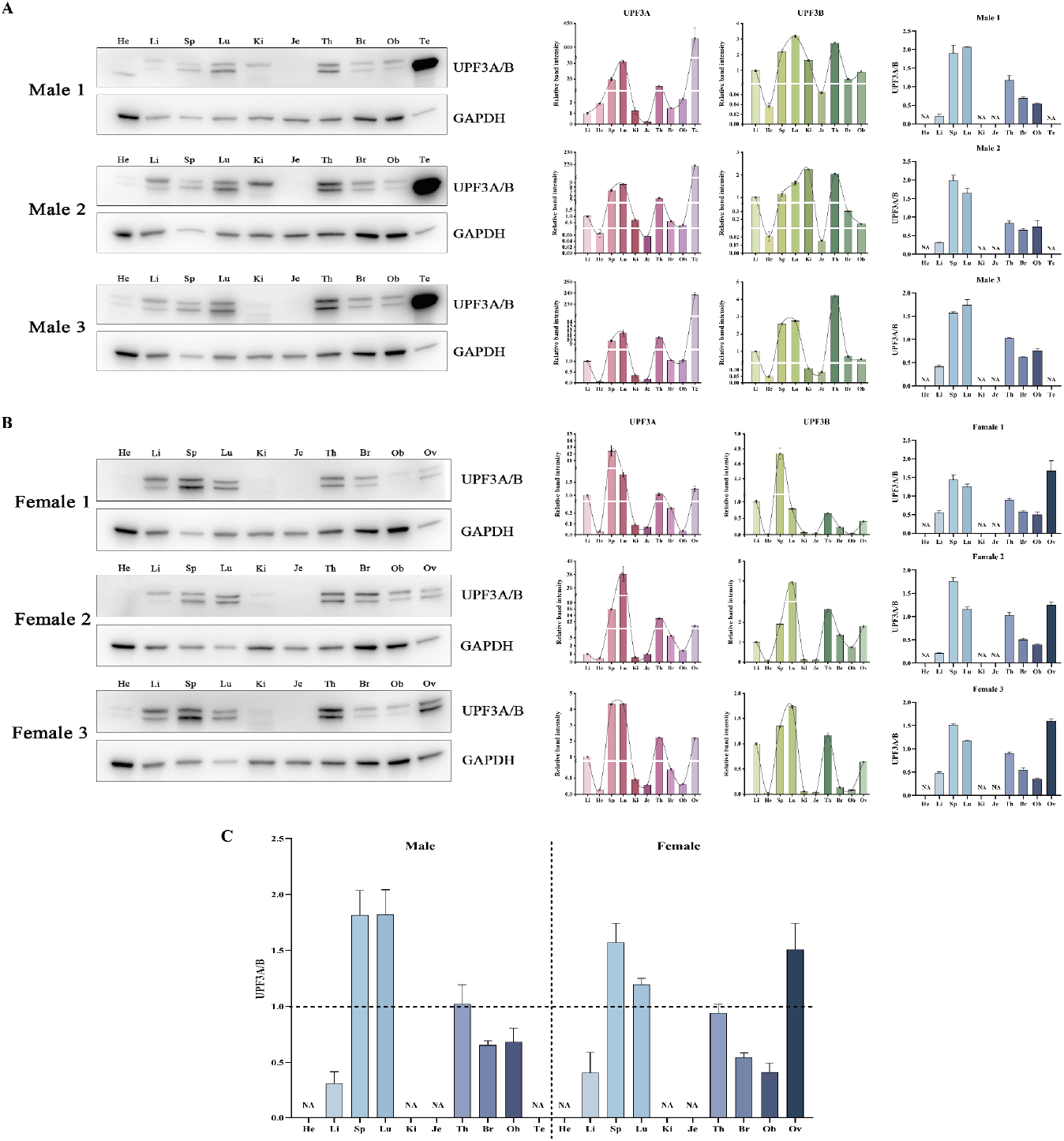
The expression of UPF3A and UPF3B proteins in tissues from C57BL/6 mice determined by Western blotting. (GAPDH was used as the loading control). **(A)** Male mice. **(B)** Female mice. **(C)** the relative protein expression of UPF3A/B. n=3. Results are represented as mean ± SD. He: heart; Li: Liver; Sp: spleen; Lu: lung; Ki: kidney; Je: jejunum; Th: thymus; Br: brain; Ob: olfactory bulb; Te: testis; Ov: ovary; NA: not available.

We further investigated the relative expression of UPF3A and UPF3B in all tissues we analyzed. We found that in spleens and lungs from both sexes, UPF3A protein levels are higher than those of UPF3B. In livers, kidneys, cerebral cortexes, and olfactory bulbs from both sexes, UPF3B is the dominant UPF3 paralog. Intriguing, we found a higher protein expression of UPF3A over UPF3B in ovaries (Figure 2). Furthermore, we went on to ask whether the expression of UPF3A and UPF3B expresses differently between sexes at given tissues. We found there was no significant difference in the protein expression of UPF3A and UPF3B in all tissues between male and female mice (Figure 3).

**Figure 3.**
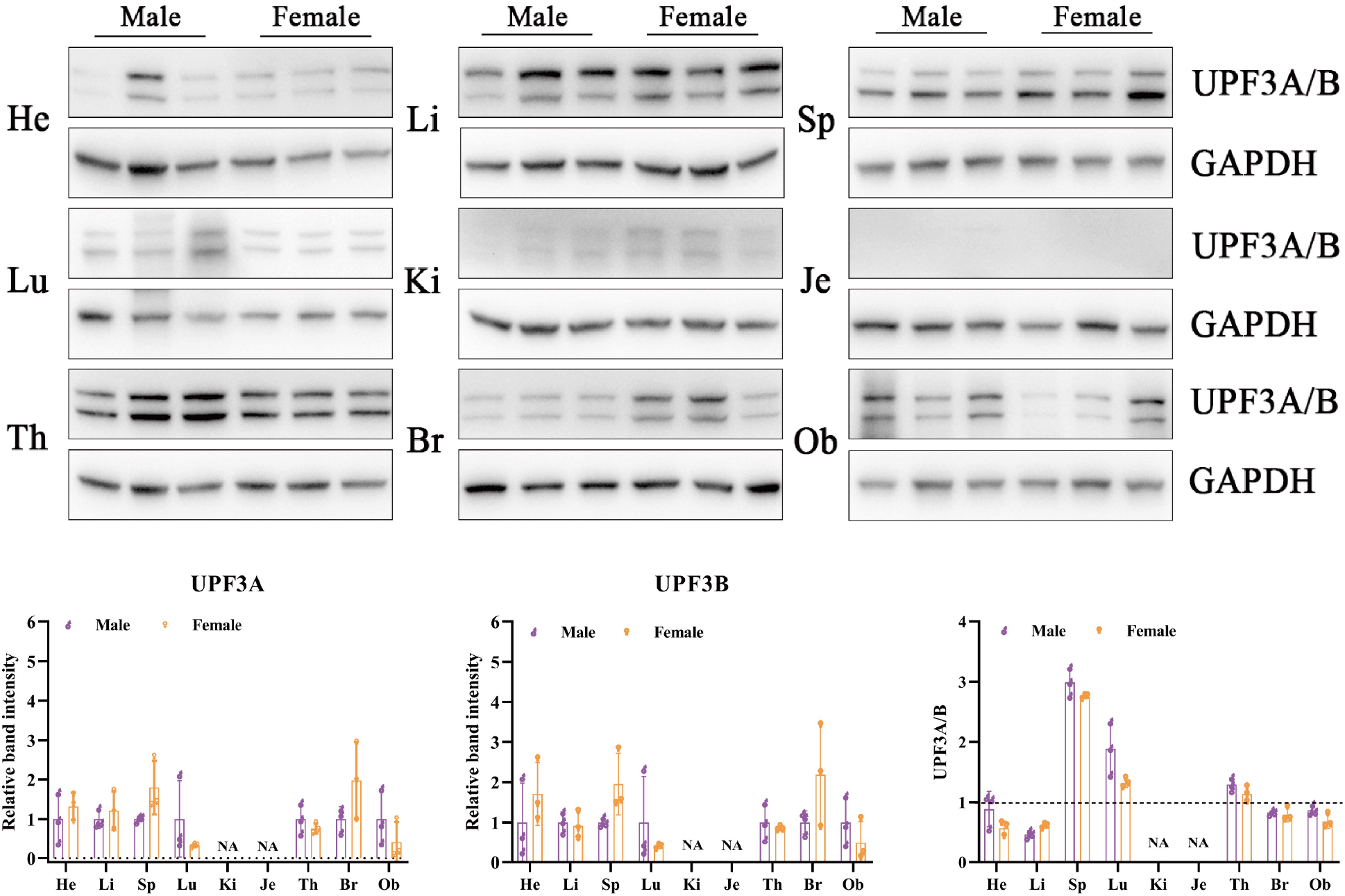
The comparison of UPF3A and UPF3B proteins in tissues from male and female mice. n=3. Results are represented as mean ± SD. He: heart; Li: Liver; Sp: spleen; Lu: lung; Ki: kidney; Je: jejunum; Th: thymus; Br: brain; Ob: olfactory bulb; Te: testis; Ov: ovary; NA: not available.

### 3.3 UPF3A is ubiquitously expressed in mouse and human tissues

To further substantiate our unexpected finding that UPF3A, as UPF3B, is ubiquitously expressed in mouse tissues, we retrieved the *Upf3a* and *Upf3b* expression data from mouse ENCODE transcriptome and used RPKM value (Reads Per Kilobase per Million mapped reads) data to reveal relative expressions of *Upf3a* and *Upf3b* in different tissues. The mouse ENCODE transcriptome dataset further confirmed that *Upf3a* is highly expressed in testis. Furthermore, the RPKM value of *Upf3a* is higher than *Upf3b* in all tissues investigated (Figure 4A).

**Figure 4.**
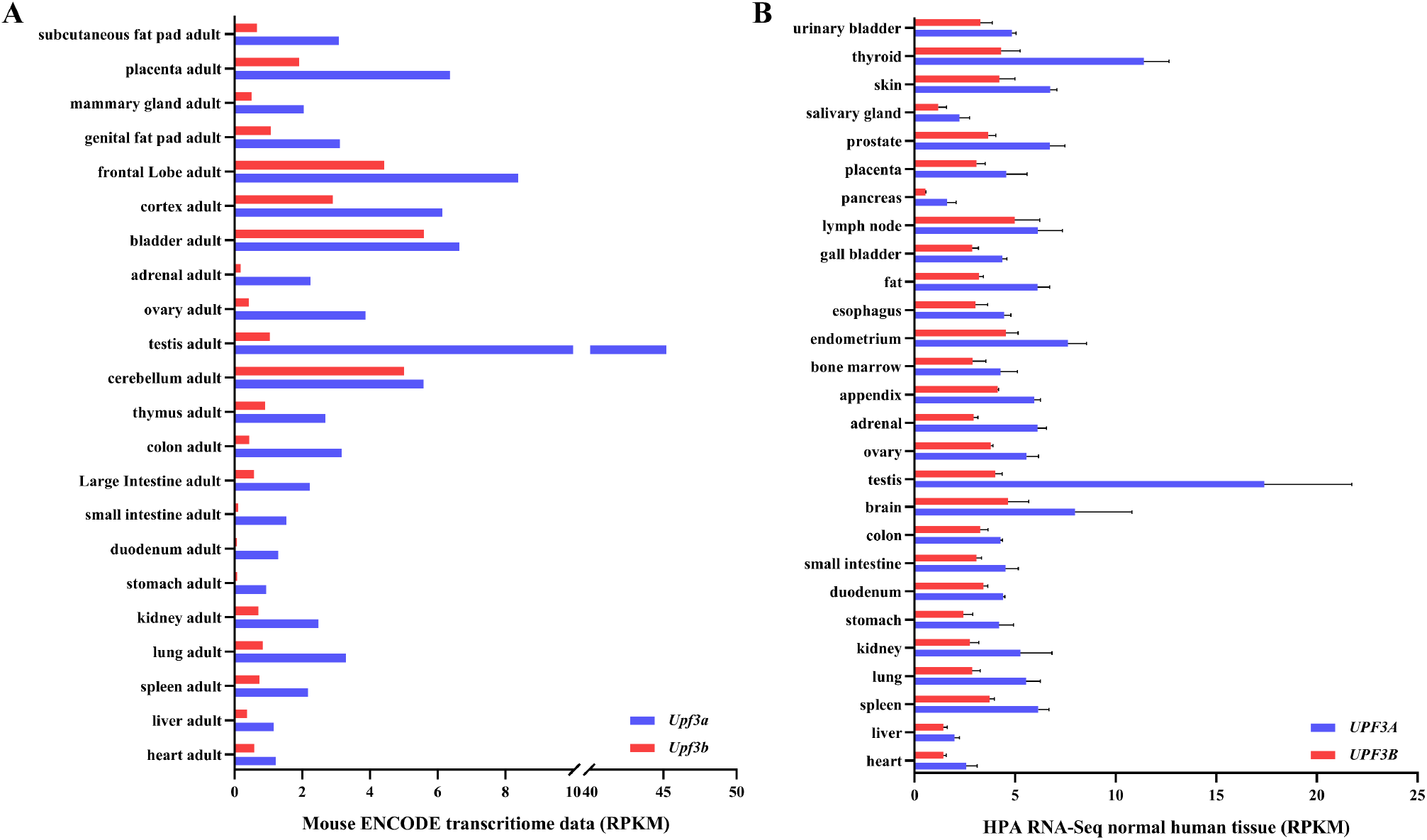
*Upf3a* and *Upf3b* mRNA levels in different organs in mouse and human. Mouse ENCODE transcriptome data **(A)** and HPA RNA-seq normal tissues **(B)** of *UPF3A* and *UPF3B* were retrieved from NCBI. RPKM, Reads Per Kilobase per Million mapped reads.

In our analysis, we further surveyed the HPA RNA-Seq data from normal human tissue, and we found that the *UPF3A* is also highest expressed in human testis (Figure 4B), implying that the expression of UPF3A is conservative. Furthermore, *UPF3A* expression is not lower than *UPF3B* expression in most tissues of human and mouse, indicating that UPF3A is ubiquitously expressed among human and mouse tissues (Figure 4A and 4B).

## Discussion

Nonsense-mediated mRNA decay is an important post-transcriptional regulatory mechanism of gene expression in eukaryotic cells (1, 3, 8-10). NMD controls the quality of transcriptome and quantity of gene transcripts, thus playing vital roles in animal development and diseases (2, 9). Although UPF3 is among the 1^st^ set of trans-regulatory NMD factors found in unicellular eukaryotic organism - yeast (13, 14), the biology of UPF3 and its vertebrate paralogs (UPF3A and UPF3B) are poorly understood. The functions of UPF3A and UPF3B are diverse and controversial (18, 22, 26-28). One of the mysterious phenomena of UPF3A and UPF3B biology is that while UPF3B is abundantly expressed in most mammalian tissues, UPF3A protein is hardly detectable (18, 20, 21). Thus, proteins of UPF3A and UPF3B could be mutually exclusively presented in most mammalian organs. We recently characterized an antibody which could detect UPF3A and UPF3B proteins simultaneously (“one tube reaction”), and found that in mouse embryonic stem cells and rib muscle derived fibroblasts, UPF3A and UPF3B are adequately expressed. This finding motivated us to investigate whether the mutually exclusive expression of UPF3A and UPF3B do exist under the physiological condition in mouse.

RT-qPCR analysis showed that mRNAs of *Upf3a* and *Upf3b* are ubiquitously presented in all tested tissues from adult male and female mice. Furthermore, the testis has the highest expression of *Upf3a*. These findings are in good constancy with previous studies (18, 20). With the antibody which could detect UPF3A and UPF3B proteins simultaneously (“one tube reaction”), we investigated the proteins of UPF3A and UPF3B in nine tissues of adult male and female mice, and compared the relative expression of UPF3A and UPF3B in these tissues. Our study confirmed that UPF3B is ubiquitously expressed, which supported the general notion that UPF3B is a major NMD related paralog of UPF3 in mammals (28). Furthermore, UPF3A is highly expressed in the testis, indicating an essential role of UPF3A in spermatogenesis (18, 20). However, the most striking finding in the current study is that UPF3A protein is also ubiquitously expressed. This finding is discrepant to most of previous studies conducted with RNA-hybridization based or antibody-based approaches (18, 20, 21). We reasoned here that our “one tube reaction” avoids several drawbacks when using immunoblotting: 1) Performance variances of individual antibody against UPF3A and UPF3B; 2) Techniques including signal detection and membrane processing. Furthermore, the ubiquitous expression nature of UPF3A is supported by publicly available RNA-seq data in mouse and human tissues. Since UPF3A knockout mice are embryonic lethal (18), the ubiquitous expression pattern of UPF3A indicates that UPF3A is essential for organ development and tissue homeostasis.

In our study, we found four types of tissue specific expression patterns related to protein levels of UPF3A and UPF3B: 1) in spleens and lungs from both sexes and ovaries from adult females, UPF3A is the major UPF3 paralog (***UPF3A^high^***); in livers, kidneys, cerebral cortexes and olfactory bulbs from both sexes, UPF3B is the dominant form of UPF3 paralog (***UPF3B^high^***) in the thymus of both sexes, UPF3A and UPF3B are comparable in protein expression (***UPF3A/B^equal^***); hearts and small intestines from both sexes express the least amount of UPF3A and UPF3B (***UPF3A/B^null^***). The biological mechanism of tissue specific expressions of UPF3A and UPF3B are worthy of further detailed characterization. Previous findings suggest that competition between UPF3B and UPF3A binding to UPF2, and thus UPF3A is destabilized when UPF3B is expressed (18, 22). Our finding here suggested that, under physiological conditions, UPF3A and UPF3B do not compete or destabilize each other. Since the co-deletion of UPF3A and UPF3B in mammalian cells showed stronger NMD inhibitory effects than UPF3A or UPF3B single deletion (16, 22, 28), our findings suggest that UPF3A and UPF3B could synergistically regulate NMD. Furthermore, the ubiquitous expression pattern of UPF3A indicates that, in addition to NMD, UPF3A may actively participate in other NMD-independent mechanisms, such as epigenetic regulation in genetic compensation, to maintain tissue homestasis (32).

In summary, in the current study, we characterized the mRNA and protein expression of UPF3A and UPF3B in ten tissues from adult male and female mice. We found that UPF3A is ubiquitously expressed, which may serve as a guidance of . Our study provides new evidence on solving the mystery of UPF3A and UPF3B biology in mammals.

## Authors’ contributions

Tangliang Li designed the experiments and supervised the project; Xin Ma performed the experiments and analyzed the data; Xin Ma and Tangliang Li wrote the manuscript, which was edited by Yan Li and Chengyan Chen. All authors read and approved the final manuscript.

## Acknowledgements

We thank all members of Li laboratory for their helpful discussions and critical comments on the manuscript. The research projects conducted in Li laboratory are supported by: Grant No. LY22C050003, from the Natural Science Foundation of Zhejiang Province, China; Grants No. KF2020005 and KF2021008 from NHC Key Laboratory of Birth Defect for Research and Prevention (Hunan Provincial Maternal and Child Health Care Hospital, Changsha, China), Grant No. 31770871 from National Natural Science Foundation of China, and Qilu Youth Scholar Startup Funding of Shandong University.

